## Supplementary figures and images for "An essential periplasmic protein coordinates lipid trafficking and is required for asymmetric polar growth in mycobacteria"

### Video Supplement 1

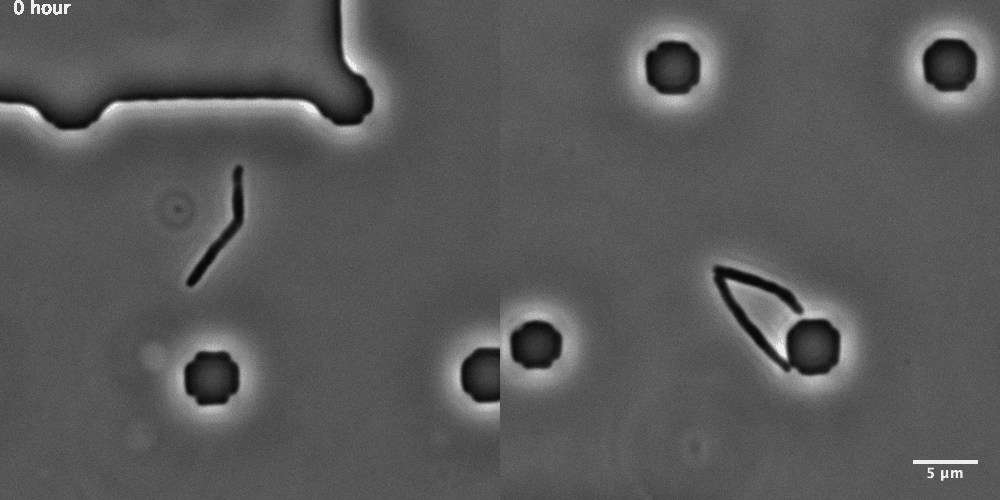
